# Evidence Accumulation Modelling Reveals that Gaussian Noise Accounts for Inhibition of Return

**DOI:** 10.1101/2020.06.21.163485

**Authors:** Tal Seidel Malkinson, Alexia Bourgeois, Nicolas Wattiez, Pierre Pouget, Paolo Bartolomeo

## Abstract

Inhibition of return (IOR) refers to the slowing of response times (RTs) for stimuli repeated at previously inspected locations, as compared with novel ones. However, the exact processing stage(s) at which IOR occurs, and its nature across different response modalities, remain debated. We tested predictions on these issues originating from the FORTIOR model (<u>fronto-parietal organization of response times in IOR</u>; Seidel Malkinson & Bartolomeo, 2018), and from evidence accumulation models. We reanalysed RT data from a target-target IOR paradigm (Bourgeois et al.,2013a, 2013b) by using a LATER-like evidence accumulation model (Carpenter & Williams, 1995), to test the predictions of FORTIOR, and specifically whether IOR could occur at sensory/attentional stages of processing, or at stages of decision and action selection. We considered the following conditions: manual or saccadic response modality, before or after TMS perturbation over four cortical regions. Results showed that the Gaussian noise parameter best explained both manual and saccadic IOR, suggesting that in both response modalities IOR may result from slower accumulation of evidence for repeated locations. Additionally, across stimulated regions, TMS affected only manual RTs, lowering them equally in the conditions with repeated targets (Return) and non-repeated targets (Non-return). Accordingly, the modelling results show that TMS stimulation did not significantly alter the pattern between model parameters, with the Gaussian noise parameter remaining the parameter best explaining the Return - Non-return RT difference. Moreover, TMS over the right intra-parietal sulcus (IPS) perturbed IOR by shortening the Return RT. When directly testing this effect by modelling the TMS impact in the Return condition, the Bayesian information criterion of the Gaussian noise parameter was the smallest, but this effect did not reach significance. These results support the hypothesis that target-target IOR is a predominantly sensory/attentional phenomenon, and may be modulated by activity in fronto-parietal networks.

## Introduction

Inhibition of return (IOR) refers to the slowing of response times (RTs) for stimuli repeated at previously inspected locations, as compared with novel ones (Berlucchi et al., 1981; Klein & Redden, 2018; Lupiáñez et al., 2006; Posner & Cohen, 1984). IOR is believed to promote our exploration of the environment by avoiding repeated scanning of previously visited locations (Klein, 1988). This phenomenon was extensively explored since its discovery in the ‘80s, but its exact nature and neural bases remain debated. Repeated peripheral events can result in faster RTs (RT facilitation) or slower RTs (IOR), depending on several variables, including the delay between the stimuli, the motor effector used (manual responses or eye saccades), and the type of visual task (detection or discrimination) (Lupiáñez, 2010). As noted by some theorists (Berlucchi,2006; Lupiáñez, 2010), this evidence challenges the view of IOR as the result of inhibiting attention from returning to a previously explored location (Klein & Redden, 2018; Posner et al., 1985). Moreover, the neural bases of IOR, as well as the exact processing stage(s) of IOR generation are currently unclear. IOR could occur at early sensory/attentional stages of processing, or at higher stages of decision and action control. Originally, Posner et al. (1985) had suggested the cause for IOR was the orienting of attention towards a location and the subsequent removal of attention from that location, to discourage attention from re-orienting back to the originally attended location. However, it was shown that this attentional account cannot fully explain IOR, and that oculomotor programming has an important contribution to IOR generation (See Klein, 2000 for a review). Converging evidence suggests that IOR could delay both motor responses and the return of exogenous attention (Lupiáñez et al., 2006).

Correspondingly, several subcortical and cortical regions were shown to be implicated in the generation of IOR. For example, neuropsychological evidence (Sapir et al., 1999) and non-human primate electrophysiology (Dorris et al., 2002) suggest an important role for the midbrain superior colliculus (SC). However, the SC is not the site where the signal reduction is generated (Dorris et al., 2002). Dorris et al. suggested that signal reduction could be potentially generated in the posterior parietal cortex (PPC). Supporting evidence for this hypothesis comes from electrophysiological studies showing that neural activity in monkey LIP was found to be reduced for already explored targets in visual search (Mirpour et al., 2009). Further evidence comes from the observation that in human patients with right hemisphere damage and visual neglect, manual IOR was reduced for right-sided, non-neglected repeated stimuli (Bartolomeo et al., 1999, 2001), where it could even revert to a “facilitation of return”, i.e. faster RTs for repeated targets (Bourgeois et al., 2012). An advanced lesion analysis in the Bourgeois et al.’s (2012) study showed that facilitation of return was found in patients with damage either to the supramarginal gyrus in the right parietal lobe, or to its connections with the ipsilateral prefrontal cortex. Importantly, however, these patients showed normal saccadic IOR. Indeed, a contribution of the frontal eye field (FEF) to IOR was also reported (Mirpour et al., 2019; Ro et al., 2003), and FEF connections with the supramarginal gyrus were shown to support visual attention shifts (Heinen et al., 2017).

One way to explore the causal role of the suggested underlying brain structures in the generation of IOR is by applying transcranial magnetic stimulation (TMS) (Satel et al., 2019). In a series of studies, Bourgeois et al. (2013a, 2013b) used repetitive TMS to test the causal role of distinct nodes of the human fronto-parietal attention networks in the generation of manual and saccadic IOR in a target-target paradigm (see Maylor & Hockey, 1985). Critically, transient interference of the intra-parietal sulcus (IPS) and temporo-parietal junction (TPJ) in either hemisphere affected IOR differently, and these effects were hemifield- and effector-dependent. Specifically, rTMS over the right hemisphere TPJ interfered with IOR only for manual responses with ipsilateral (right) targets, consistent with data obtained in right-brain damaged patients with neglect (Bourgeois et al., 2012). TMS over the right hemisphere IPS perturbed manual IOR for targets on both sides of the screen, but saccadic IOR was affected only for contralateral (left) targets (Bourgeois et al.,2013b). Left hemisphere stimulation did not affect IOR (Bourgeois et al., 2013a). More recently, Seidel Malkinson & Bartolomeo (2018) proposed a theoretical model labeled FORTIOR (Frontoparietal Organization for Response Times in IOR), to explain the cortical basis of IOR generation in target-target detection paradigms, and to account for the complex pattern of interference produced by TMS stimulation. Based on the known architecture of frontoparietal cortical networks and on their anatomical and functional asymmetries (Bartolomeo & Seidel Malkinson, 2019), FORTIOR postulates that both manual and saccadic IOR arise from a noise-increasing reverberation of activity within priority maps of the frontoparietal circuit linking frontal eye field (FEF) and intraparietal sulcus (IPS). The noisier saliency signal is then forwarded to the manual and saccadic motor systems, and results in IOR. Differences between the readout capacities of the manual and saccadic effector systems when reading the output of the FEF-IPS circuit may then lead to the dissociations between saccadic and manual IOR. Klein and Redden (2018) suggested a different view, and proposed the existence of two types of IOR: an ‘input’ form causing IOR by biasing perception, and an ‘output’ form causing IOR by biasing action. Specifically, Klein and Redden suggested that input-IOR is generated when the reflexive oculomotor system is actively suppressed (ie, when participants must maintain fixation and respond manually), and results from modulation of activity in early sensory pathways in retinotopic coordinates, rather than from response outputs. Output-IOR occurs when the reflexive oculomotor system is not suppressed, and involves projections from the SC to cortical areas (i.e. FEF and PPC). Output IOR is spatiotopic and is independent of the response modality. Therefore, according to this account, manual IOR (while maintaining fixation) and saccadic IOR reflect different signal reduction processes and implicate different underlying neural mechanisms.

Accumulator models, which explain the decision to respond as an accumulation of evidence to a response threshold (see also reviews by Duncan Luce, 1986; Ratcliff & Smith, 2004), can provide an important contribution to our understanding of IOR. Both behavioural and neurophysiological evidence suggest that the accumulation rate is affected by the stimulus quality and that the evidence criterion is modulated by the prior probability of a particular response (Gold & Shadlen, 2007; Pouget et al., 2011). In the case of IOR, increased RTs for repeated targets might be explained by a variety of model parameters (Satel et al., 2019), such as sensory-level effects (e.g., slower rate of accumulation or decreased noise), or a higher decision-level effect (higher baseline or decision threshold, jointly referred to here as threshold). The identification of the relevant parameters may thus help determine the processes implicated in IOR. Presently, only two studies used this approach to explore IOR. The first one modelled RT data from two experiments, in which participants generated sequences of three saccades in response to a peripheral or a central cue, generating saccadic IOR for movements to the immediately preceding fixated location (Ludwig et al., 2009). Saccadic IOR was best accounted for by a change in the accumulation rate. Ludwig et al. (2009) concluded that an attenuated sensory response to stimulation at a recently fixated location might be responsible for a lower accumulation rate for a response to that location. This finding was the first to propose a plausible explanation for a reduced accumulation rate under peripheral cueing conditions. However, the authors did not extend their finding to manual responses, and therefore could not provide evidence of such an account for the data produced with a non-specific effector (see also Rafal et al., 1994; Taylor & Klein, 2000).

In the second study, MacInnes (2017) modelled data from two experiments testing the gradient of IOR with random, continuous cue-target Euclidean distance and cue-target onset asynchrony. Although the behavioral RTs differed across response modalities, best-fit models were shown to have similar model parameters for the gradient of IOR for both effector types, suggesting similar underlying mechanisms for saccadic and manual IOR. Contrary to Ludwig et al. (2009), MacInnes (2017) found that the model parameter best explaining location-associated changes in RT was the variance of accumulation starting point for both manual and saccadic responses. When the spatial distance between the location of the cue and the target increases, the variability in the starting point of information accumulation also increases. This, on average, shifts the accumulation closer to the decision threshold and thus decreases RTs. This result is consistent with neural results where changes in resting state activation levels in SC (Dorris & Munoz, 1998) are dependent on target location probabilities. The conflicting results of IOR-associated model parameters in the Ludwig et al. (2009) and MacInnes (2017) studies may result from differences in the experimental paradigm.

To address these issues, here we used an evidence accumulation model inspired by the LATER model (linear approach to threshold with ergodic rate; Carpenter & Williams, 1995; see also for review Noorani, 2014; Schall, 2019). Conceptually minimalist and mathematically tractable, this is a simple model for RTs. In this model, information linearly accumulates until reaching a criterion. The rate of accumulation varies between responses in a Gaussian manner and a uniform distribution of starting points. A simple and formal view of this model is illustrated in Figure 3 below, which shows what may be regarded as the ‘decision period’ of a fixation period prior to saccadic or manual initiation response. An additional, constant non-decisional latency that includes afferent and efferent delays associated with stimulus encoding and peripheral motor delays is also added to ‘the decision period’ (Brown & Heathcote, 2008; Pouget et al., 2009). In LATER, a decision signal starts from a starting point S_0_ and rises toward a threshold θ. Once the signal reaches the threshold, a decision is made for a particular action. The rate at which the decision signal rises varies randomly from trial to trial, but the mean rate of rise is constant and denoted by the parameter μ. The standard deviation of this variation in rate of rise is given by the parameter σ (see Figure 3). Together, these assumptions of variability in starting point and accumulation rate are sufficient to produce latency distributions that have the appropriate typical right-skewed shape.

In the present study, we used an evidence accumulation model to further explore the mechanism of IOR, the dissociation between saccadic and manual IOR, and the nature of TMS-induced IOR perturbations. Specifically, we had three aims: 1. Determine if IOR RT-distributions could be best accounted for by sensory parameters (rate of accumulation and noise), or by decisional processes (decision threshold); 2. Test whether the same model parameters can account for RT-distributions for manual and saccadic IOR, and thus reflect a common mechanism; 3. Explore if TMS-induced IOR perturbation operates at a sensory or a decisional level.

## Methods

### Experimental data

The RT data used for modelling was taken from the studies by Bourgeois et al. (2013a, 2013b), who used repetitive TMS to assess the causal role of distinct nodes of the human frontoparietal networks in attentional orienting. In these studies, participants performed a target-target paradigm (see Maylor & Hockey, 1985). Four black peripheral circles surrounding a circle placed at fixation were displayed (see Fig. 1). Participants had to respond as fast as possible to one of the peripheral circles becoming white, either by pressing a key while maintaining fixation, or, in a different session, by making a saccade towards the target. We compared RTs for targets appearing at the same location of the preceding one (Return trials) with RTs for targets appearing at new locations at the horizontally and vertically opposite side of the display (ie, a bottom-left target after a top-right target; Non-return trials). Responses to right-sided and left-sided targets were pooled together inside each return category to maximize modelling robustness. The TMS-stimulated brain nodes were the intraparietal sulcus (IPS) and the temporo-parietal junction (TPJ) in the left or in the right hemisphere. We modelled RT data from 32 participants (8 in each TMS-stimulation group).

**Figure 1.**
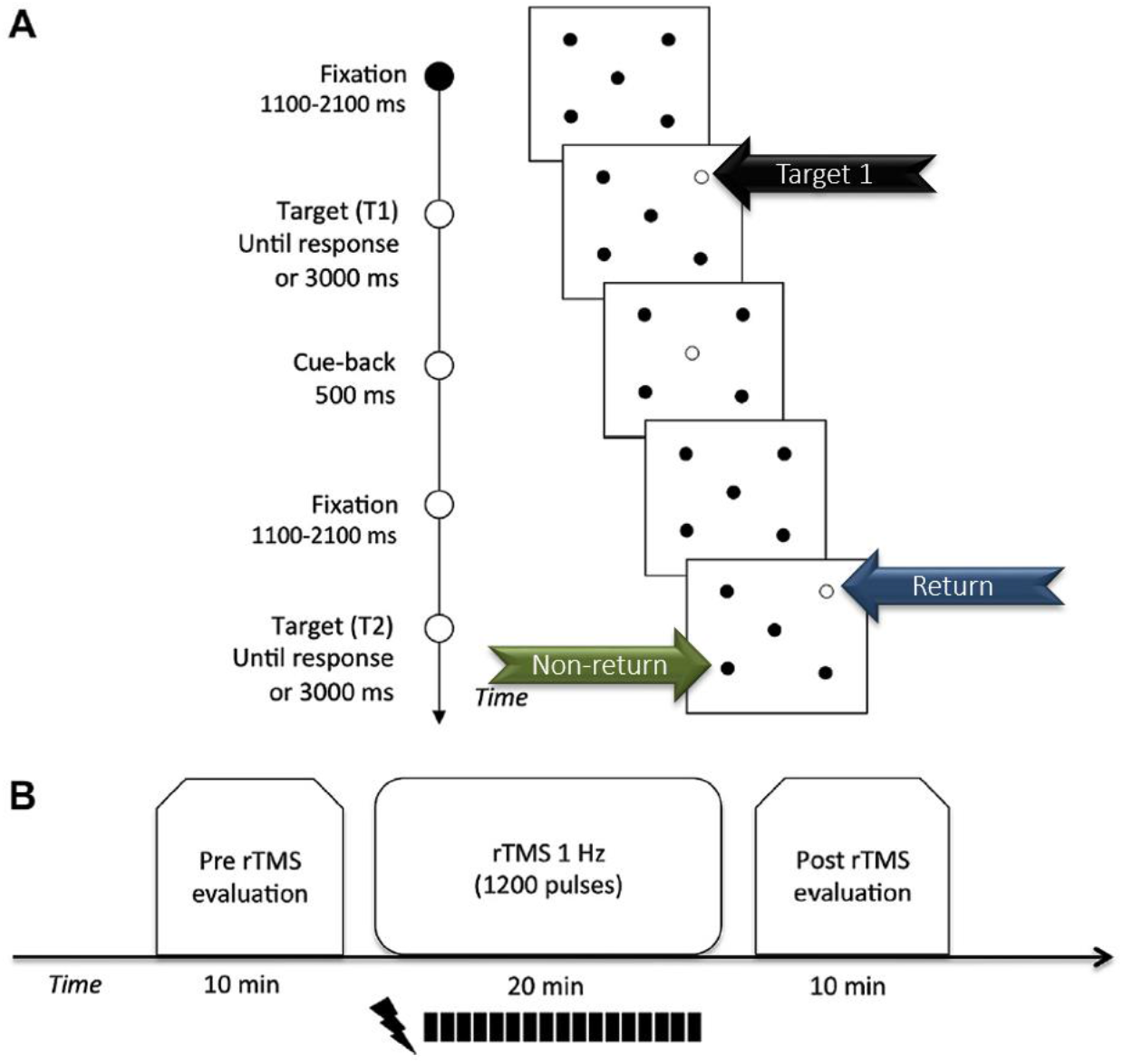
Target-Target detection task in Bourgeois et al., 2013a, 2013b. A. Trial sequence and timing. In the manual task, participants maintained fixation and manually detected the appearance of peripheral targets. In the saccadic task, participants moved their gaze to peripheral targets and back to the center upon cue-back appearance. B. Timeline of the behavioral and rTMS conditions. Two runs of each task (manual and saccadic) were performed for each participant in two different sessions. A 10 min run was performed immediately before (pre-TMS) and the other one immediately after (postTMS) repetitive TMS stimulation. Repetitive TMS patterns consisted of 1200 TMS pulses applied at 1 Hz with an inter-pulse interval of 1 sec (for a total of 20 min). The data analyzed in this study consisted of only the Targets repeating in the same exact location (Return trials, blue arrow) and targets repeating in the farthest location (Non-return trials; green arrow) from the previous target (Target 1; black arrow). Adapted from Bourgeois et al., 2013.

### Model fitting

The linear ballistic accumulator model assumes between-trial noise such that the rate on a simulated trial is drawn from a normal distribution with mean given by the mean rate and standard deviation σ (Brown and Heathcote, 2008). Its activity (dago) is governed by a stochastic differential equation:

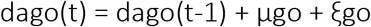

The unit is defined by two parameters: the mean growth rate (μgo) and a Gaussian noise term (σgo) with a mean of zero variance of ξgo, where a represents the activity of the unit. The race finished when the unit crossed the threshold (θ), within a limit of 3000 ms. If unit activity is negative during the race, the activity was reset to zero at this point (non-physiologic value). An additional parameter, Dgo, was added to the model to take into account the stimulus encoding that occurred in the go unit. Dgo was set to 100 ms. μgo, σgo and θ are the unconstrained parameters of the model.

The observed latency distributions for correct responses as the latency distributions generated by the model were binned into 5 quantiles, defined by the following boundaries: {0, 0.2, 0.4, 0.6, 0.8, 1.0}. A local chi-square was computed at each bin and a general chi-square was obtained by summing each local chi-square. To find the best parameters for each latency distribution, we minimized the general chi-square by using a minimization function (patternsearch from the global optimization toolbox of Matlab). Patternsearch looks for a minimum based on an adaptive mesh that is aligned with the coordinate directions. Because minimization functions are sensitive to their starting point, we ran pattern search from 50 randomly chosen starting points. Finally, to avoid a local minimum, we again started patternsearch from 200 new start points. These starting points were determined by the best parameters from the first run and were defined as follows: best parameters ± 0.01 units and best parameters ± 0.02 units. After applying this procedure to each distribution of the Return and the Non Return conditions, we ran a new procedure in order to determine which of the μ, σ or θ parameter could explain the differences observed in the Return and the Non-Return latency distributions. To do this, we fixed two parameters of the Return condition on latency distribution of the Non-return condition and let one parameter free. As in the first procedure, pattern search was run from 50 randomly chosen starting points and then from 200 new start points. All these procedures were performed on a supercomputer cluster (NEC, 40 nodes, 28 cpu Intel Xeon E5-2680 V4 2.4 GHz/node, 128 Go RAM/node).

The analysis code can be obtained by request to the corresponding author.

### Statistical analysis

Following <u>Ratcliff and Tuerlinckx (2002)</u>, optimized fit of each model can be minimized using (χ^2^) statistics (Pouget et al. 2011). However, to account for the number of parameters as well as the variance, here we used Bayesian information criterion (BIC) values that were computed for each model using Matlab (MATLAB, 2017) and served for model estimation and comparison. RTs and BIC values were statistically compared using the JASP software (Jasp Team, 2020, JASP (Version 0.12)[Computer software]). We modelled RT data from 32 participants (8 in each TMS-stimulation group). In order to keep inter- and intra-subjects RT variability and in accordance with the procedure in Bourgeois et al., (2013a, 2013b), trials with RTs shorter than 100ms (anticipations), as well as trials with RTs slower than 2.5 SD were excluded. For RT analysis, these same trial-selection criteria across the 32 participants were used. However, the present trial selection procedure differs from the one originally used by Bourgeois et al. (2013a, 2013b), in that it pooled together data for right-sided and left-sided targets in order to maximize the number of modelled trials. When possible, repeated measures ANOVAs (with Greenhouse-Geisser correction when necessary) and Holm-corrected post hoc tests were used for RT and model parameter analyses. For the right IPS condition, manual RT data did not deviate significantly from a normal distribution (Shapiro Wilk test, Return preTMS: W=0.950, *p*=0.71; Return postTMS: W=0.947, *p*=0.68; Non-return preTMS: W=0.972, *p*=0.91; Non-return preTMS: W=0.917, *p*=0.41). Similarly, BIC values did not deviate significantly from a normal distribution (Shapiro Wilk test, Return μ-free, σ-free and θ-free models: W=0.960, *p*=0.81; W=0.930, *p*=0.51; W=0.962, *p*=0.83; respectively; Non-return μ-free, σ-free and θ-free models: W=0.948, *p*=0.67; W=0.933, *p*=0.54; W=0.975, *p*=0.93, respectively). Therefore, repeated measures ANOVA was used to test the main effects of Return/Non-return and TMS and their interaction on RTs and BIC values.

No part of the study procedures and analyses were pre-registered prior to the research being conducted.

## Results

Across subjects, after excluding anticipatory responses and responses slower than 2.5 SD, the median number of included trials for manual responses in the Return and Non-return conditions was 30 (range, 14-49) and 33 for saccadic responses in Return and Non-return conditions (range, 20-72).

Fig. 2 shows the mean saccade latencies and manual response times across subjects, for each of the conditions of Return vs Non-return, preTMS and postTMS. A number of strong trends emerge. First, a three factor repeated-measures ANOVA was performed to assess the reliability of the Return and the effect of TMS across both manual and saccadic modalities. The main effect of Return versus Non-return was significant (F_(1,31)_=16.67, *p*<0.001, **η**^2^=0.019; see figure 2). There were also significant main effects of Modality (F_(1,31)_=12.13, *p*<0.001, **η**^2^=0.19) and of TMS (F_(1,31)_=13.48, *p*<0.001, **η**^2^=0.028). There was no interaction between Return and modality, suggesting that IOR was independent of response modality. However, the interaction between Return and TMS was significant (F_(1,31)_=5.76, *p*=0.023, **η**^2^=0.004), resulting from a significant TMS effect on the Return condition and from a significant RT difference between preTMS Return and pre and post TMS Non-Return conditions (all three post-hoc tests: *p*<0.001), similarly to the original studies by Bourgeois et al. (2013a, 2013b). Additionally, TMS and Response modality interacted (F(1,31)=21.39, *p*<0.001, **η**^2^=0.047), because only manual responses were affected by TMS (Post hoc tests: Saccadic response preTMS VS postTMS: *p*=0.46; Manual response preTMS VS postTMS: *p*<0.001). Importantly, the simple main effect of Return VS Non-Return in both manual and saccadic preTMS conditions was significant (*p*=0.002 and *p*<0.001, respectively), demonstrating the existence of a significant IOR effect prior to the TMS perturbation in both modalities.

**Figure 2:**
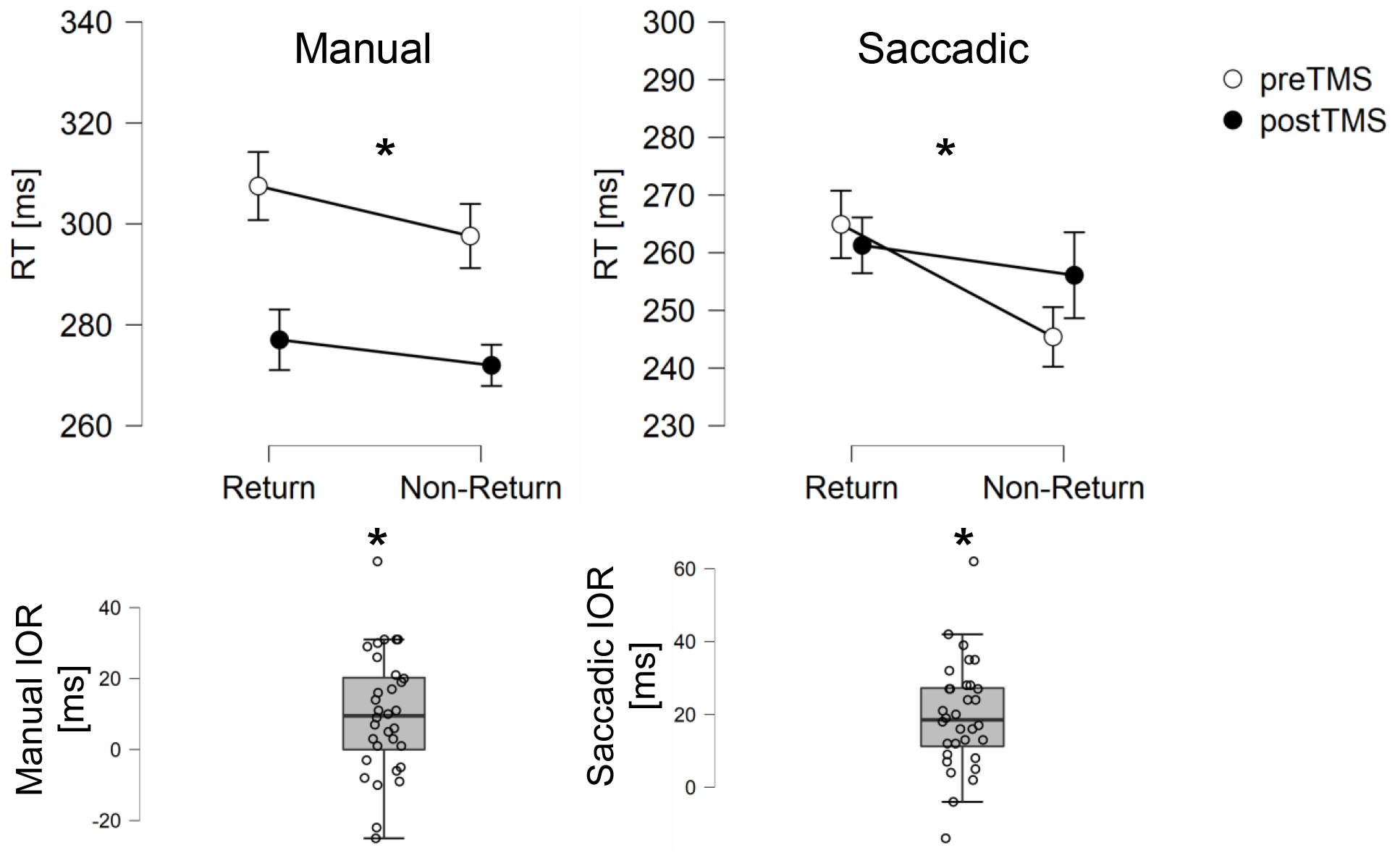
Response time inhibition of return (IOR) effect. Top: Manual response time (RT, left) and Saccadic RT (right) are longer in Return trials than in Non-return trials (Return main effect, p<0.001), preTMS perturbation (white circles) and postTMS perturbation (black circles). TMS perturbation affected manual responses but not saccadic responses (simple manual TMS effect: p<0.001; simple saccadic TMS effect: p=0.44). Error bars represent SE. Bottom: Box plots of the manual (left) and saccadic (right) preTMS IOR effects across participants. Manual and saccadic IOR effects were significant (*p*=0.002; *p*<0.001; respectively).

Figure 3 shows illustrative examples of reciprobit plots that correspond to the log-log cumulative probability distribution of saccadic or manual RTs for Return and Non-return conditions. The figure illustrates examples of single sessions showing RT variations best fitted with a single parameter adjustment (μ, σ and θ). If IOR is mediated by a change in the underlying rate of accumulation, the rate should be lower for Return responses compared to Non-return responses. If IOR reflects the need for more evidence, the threshold should be elevated for Return responses. The mean accumulation rate for Return responses is, on average, lower than that for Non-return responses.

**Figure 3.**
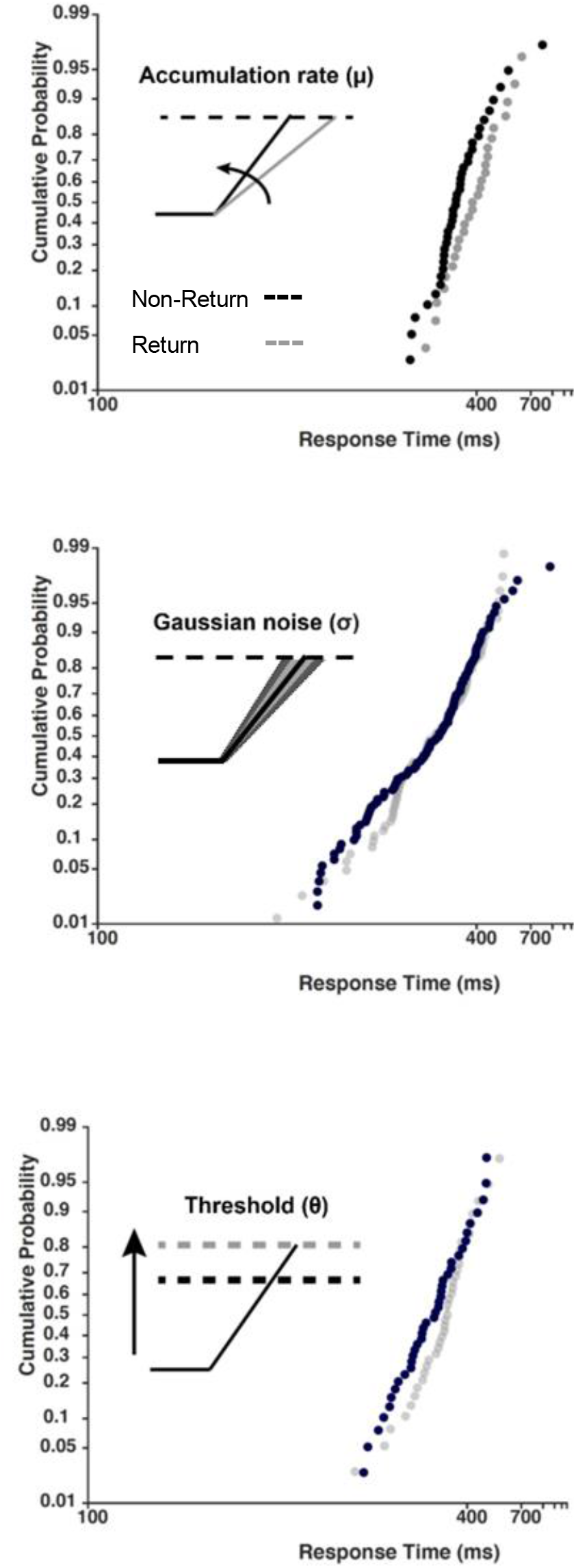
Illustrative examples of reciprobit RT distributions for Return (grey dots) and Non-return (black dots) trials. Inset illustrates the model parameter accounting for response time differences due to inhibition of return (IOR). Race to threshold between two independent accumulators, each subject to Gaussian variability in the accumulation rate and uniform variability in the starting point of accumulation. The accumulator coding the Return condition is shown by the grey line. The black accumulators code the Non-return condition. (Top panel) The target is at a Return location, causing a reduction in the accumulation rate of the associated movement program. As a result, the threshold is crossed later. (Middle panel) The target is at a return location (grey dots), resulting in a selective increased variability in the accumulation rate of the accumulator coding the target and a delay in the time at which the threshold is reached. (Lower panel) The target is at a return location, resulting in a selective increase in the evidence criterion for the accumulator coding the target and a delay in the time

### The Gaussian noise parameter σ explains manual and saccadic IOR

RTs of Return trials and Non-return trials were separately fit with accumulator models with 3 free parameters. The resulting Return condition model parameters were used to constrain the Nonreturn model fit. BIC values were computed for each of the Non-return single-free parameter models, allowing to explore which parameter best accounted for the differences between Return and Non-return trials caused by IOR. This procedure was performed separately for manual and saccadic RTs, preTMS and postTMS.

In order to test which parameter: μ, σ or θ, *explained* best the differences between Return and Non-return conditions across modalities and TMS conditions, we conducted a 3-way-repeated measures ANOVA with Free model parameter (three levels: μ-free, σ-free or θ-free models), Modality (Manual or Saccadic) and TMS (preTMS or postTMS) as factors. A significant main Free model parameter effect (F_(1.188,36.82)_=10.43, *p*=0.002, η^2^=0.021; see Figure 4), resulting from significant differences between the BIC of the σ-free and the and μ-free and θ-free models (Post Hoc comparisons: *p*=0.006). There was also a significant main effect of Modality (F_(1,31)_=11.94, *p*=0.002, η^2^=0.094), in which saccadic BIC values were smaller than manual BIC values. There was no significant TMS effect and no significant interactions between the factors. These results suggest that variability in the rate of accumulation best explains RT differences between Return and Non-return trials across both modalities, and thus support a perceptual/attentional account of IOR in both manual and saccadic modalities. Additionally, these results show that TMS stimulation did not modulate the inter-parameter BIC pattern, leaving the σ free-parameter the best accounting parameter for Return vs Non-return differences.

**Figure 4.**
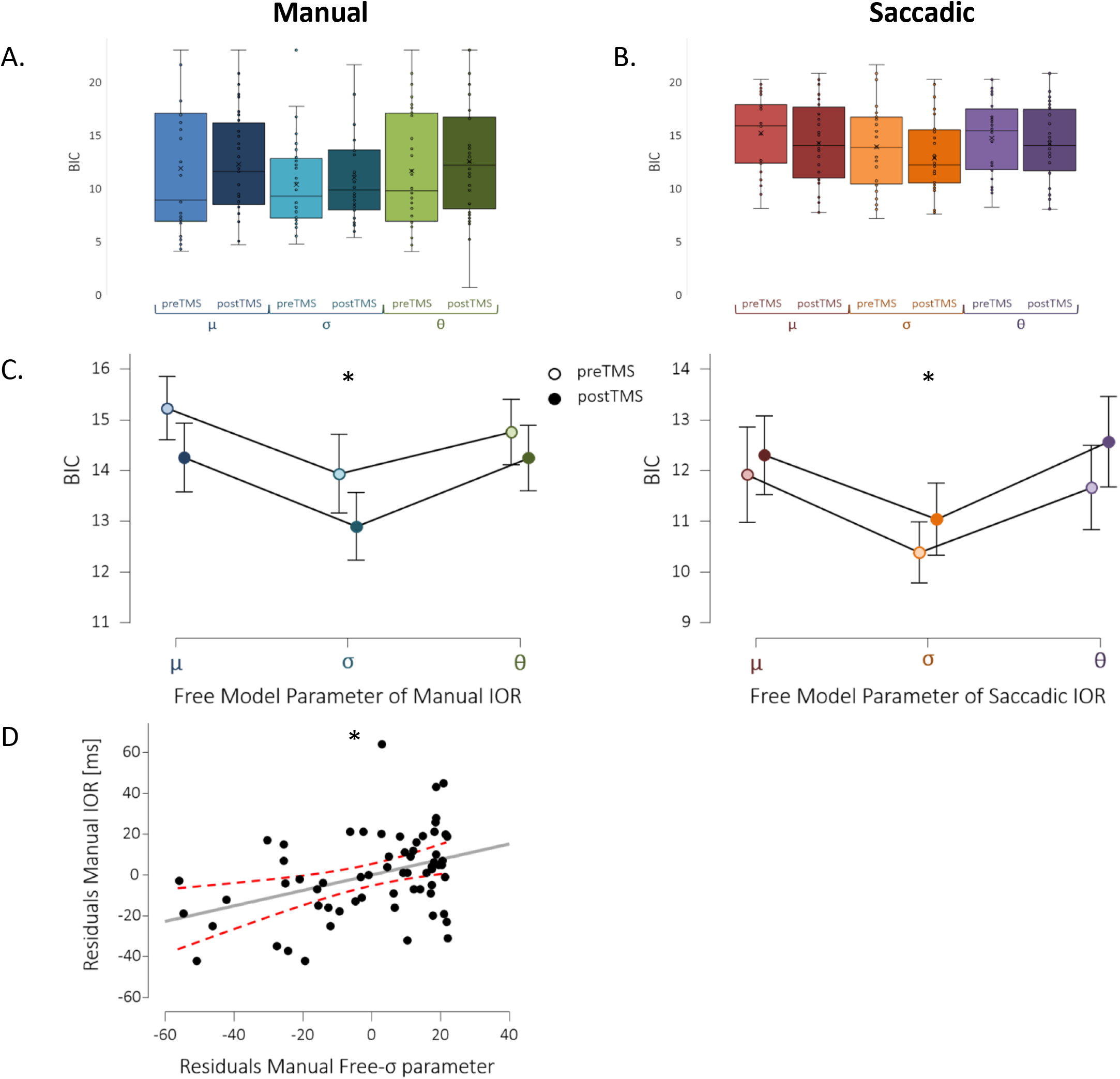
Gaussian noise parameter best explains IOR effect. A. Boxplots of BIC values for μ, σ & θ free models of manual IOR, preTMS & postTMS. B. Boxplots of BIC values for μ, σ & θ free models of saccadic IOR, preTMS & postTMS. C. Gaussian noise parameter (σ) is significantly the smallest in in manual (left panel) & saccadic modalities (right panel), preTMS (light symbols) & postTMS (dark full symbols; Free model parameter main effect, *p*=0.002; Post hoc tests: *p*<0.001). Error bars represent SE. D. A partial plot showing the relationship between the manual free-σ parameter & manual IOR, accounting for the other regression model predictors (intercept & TMS). Dashed red lines indicated 95% confidence interval. The free-σ parameter significantly predicted manual IOR (σ coefficient *p*=0.001).

To further explore the relationship between the Gaussian noise parameter and IOR we performed a multiple linear regression to predict IOR based on the σ parameter and TMS, separately in the manual and saccadic modalities. In the manual modality, a significant regression was found (F_(2,61)_=6.044, *p*=0.004), with an R^2^ of 0.138, but only the σ predictor was significant (σ coefficient: *p*=0.001; TMS coefficient: *p*=0.48). Predicted manual IOR was equal to −22.89+0.39*σ+3.49*TMS, where σ was measured in arbitrary units, and TMS was coded as 1=preTMS and 0=postTMS. Comparing a regression model including only the intercept and TMS to a model also including theσ parameter showed that the addition of the σ parameter increased the R^2^ by 0.152. In the saccadic modality, this regression was not significant (F_(2,61)_=2.81, *p*=0.068). However, a model containing only the intercept and the TMS predictor was significant (F_(2,61)_=5.052, *p*=0.028), with an R^2^ of 0.075, and a significant TMS coefficient (*p*=0.036). Predicted saccadic IOR was equal to 4.75+14.75*TMS, where TMS was coded as 1=preTMS and 0=postTMS. Comparing a regression model including only the intercept and TMS to a model also including the σ parameter showed that the addition of the σ parameter increased the R^2^ by 0.009. Thus, the σ parameter significantly predicted the IOR in the manual modality but not in the saccadic modality. This might have resulted from the limited sampling rate in the saccadic RT measurement, restricting the variability of saccadic RTs (see Discussion).

### TMS perturbation over right IPS is explained by the slope parameter

In the original studies by Bourgeois et al. (Bourgeois et al., 2013a, 2013b) TMS stimulation over the right IPS perturbed manual IOR. Analyzing thes σ manual RT data using a 2-way repeated measures Anova with Return (Return or Non-return) and TMS (preTMS or postTMS) as factors, showed a significant TMS effect (F_(1,7)_=10.74, *p*=0.014, η^2^=0.489; see Figure 5) and a TMS x Return significant interaction (F_(1,7)_=5.68, *p*=0.049, η^2^=0.056). This interaction resulted from a significant TMS effect in the Return condition (Post Hoc comparisons: Return preTMS VS post TMS *p*=0.016; Non-return preTMS VS post TMS *p*=0.16). The main effect of TMS was also evident

**Figure 5:**
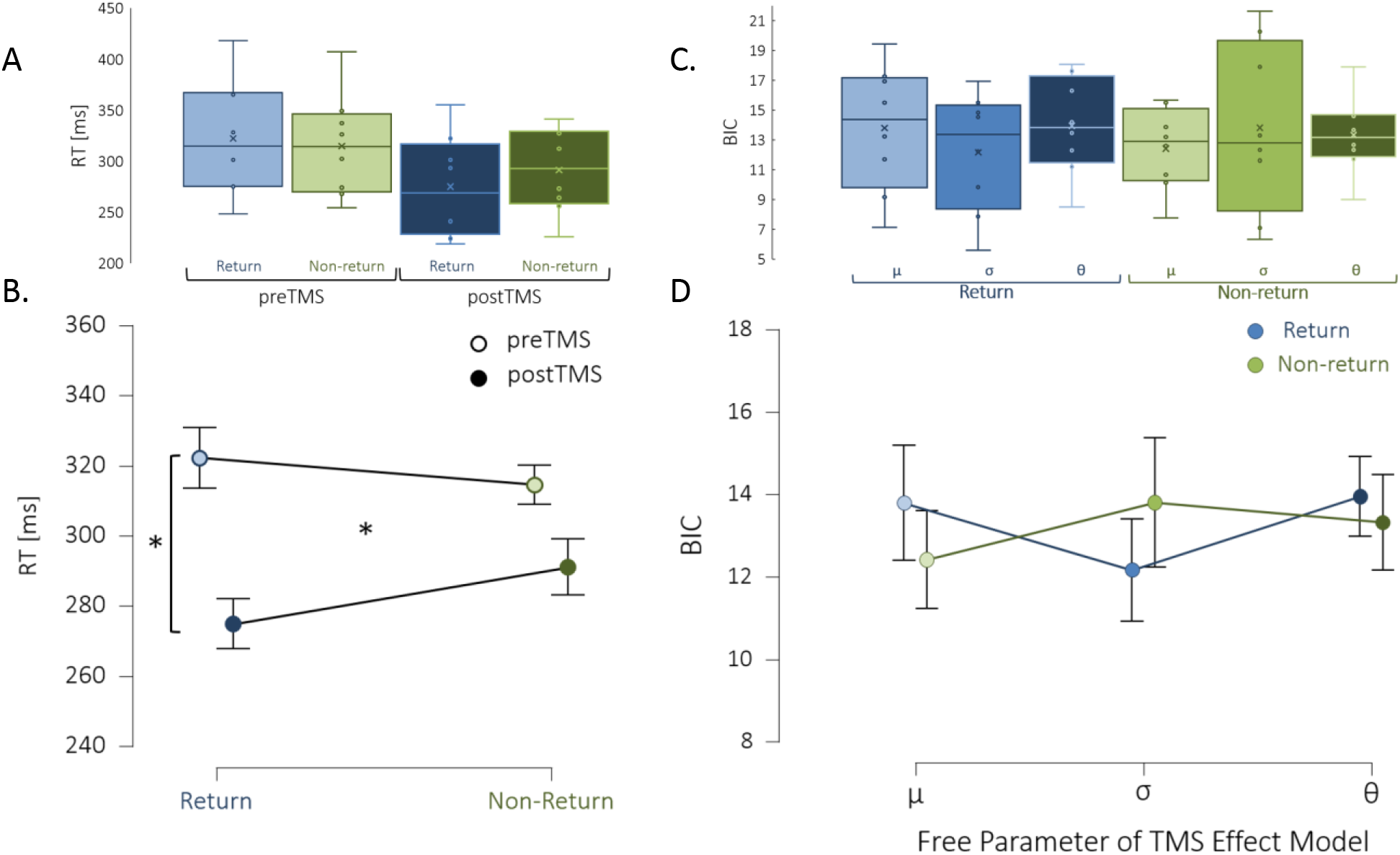
TMS perturbation over right IPS affects manual IOR. A. Boxplots of the manual RTs in the Return (blue) & Nonreturn (green) conditions, preTMS (right) & postTMS (left) of the 8 participants in the right IPS TMS group. B. Manual IOR effect preTMS stimulation (light circles) and postTMS IOR (dark circles). RT in the Return condition significantly changed following TMS stimulation (2-way repeated measures Anova: TMS x Return interaction - F_(1,7)_=5.68, *p*=0.049, η^2^=0.056; Post hoc test: *p*=0.016), causing a TMS-induced IOR effect perturbation. C. Boxplots of the BIC values of the μ-free, σ-free & θ-free TMS-models in the Return (blue) & Non-return (green) conditions. D. BIC values of TMS effect models in the Return condition (blue circles) and Non-return condition (green circles). In the Return condition, Gaussian noise parameter was the smallest and in the Non-return condition Accumulation rate parameter was smallest, but these trends did not reach significance. Error bars represent SE.

Next, we wanted to explore the origin of these observed behavioral effects of TMS perturbation over the right IPS. To this end, independently for Return and Non-return conditions, RTs of preTMS and postTMS trials were separately fit with accumulator models with 3 free parameters. The resulting preTMS model parameters were used to constrain the postTMS model fit. BIC values were computed for each of the postTMS single-free parameter models, allowing to explore which parameter best accounted for TMS induced perturbation of IOR observed behaviorally. In the Return condition, the σ-free BIC was the lowest (Return BIC Mean±SD: σ-free BIC 12.17±4.03; μ-free BIC 13.79±4.27; θ-free BIC 13.95±3.29), suggesting that Gaussian noise parameter explained the significant TMS-induced RT effect in the Return condition. In the Non-return condition, a different tendency was observed. Here, the μ-free BIC was the lowest and the σ-free BIC was the highest (Non-return BIC Mean±SD: μ-free BIC 12.42±2.75; θ-free BIC 13.36±2.60; σ-free BIC 13.80±5.71). To test the significance of these effects, we conducted a 2-way repeated measures Anova with Free model parameter (three levels: μ-free, σ-free or θ-free models) and Return (Return or Non-return) as factors, but this did not reveal significant effects (see Figure 5).

## Discussion

Does IOR attenuate processes at the sensory/attentional level, or does it influence motor programming? Behavioral evidence suggests that the sensory attenuation of stimulus-related responses may be automatic and non-motor, because it occurs independently of whether an eye or hand response is required, or whether the validity of the cues elicits reduced or increased reaction times (see, e.g., Lupiáñez et al., 2004; Posner & Cohen, 1984). Studies on the neurophysiological substrate of IOR in humans and non-human primates have led to similar conclusions. Reduction of sensory cortical responses has been observed using an event-related potential IOR task (McDonald et al., 1999) and in response to repeated stimuli using a non-RT, delayed match to sample tasks both in the PPC (Steinmetz et al., 1994; Steinmetz & Constantinidis,1995) and in the inferior temporal cortex (Miller et al., 1991). This suggests that it is a ubiquitous response property throughout a variety of brain areas. In the oculomotor sub-cortical system and in SC in particular, neuronal recordings have shown correlations between reduction of the early sensory neuronal activity in SC and IOR behavior across experimental sessions and on a trial by trial basis within a single session (Dorris et al., 2002). All these reduced stimulus-related responses are compatible with a sensory-attentional account of IOR. Despite all these results, there is still debate on whether certain additional motor components of IOR remained valid or not (Ivanoff & Klein, 2001; Taylor & Klein, 2000). Mechanistically, a signal-to-noise ratio reduction of the sensory response applied to previously attended objects or regions of a scene is one way to explain IOR. This process would ensure that an initially salient region of the scene might not continue to have this status in time. Neurophysiologically, across multiple primary sensory areas, when a sudden stimulus appears at a position outside the receptive field (RF), lateral inhibition dominates the responses inside the RF (Pouget et al. 2005; Schall and Hanes 1993). When sequential stimulations are presented, RF responses are modulated by lateral inhibition. More precisely, the lateral inhibition following the first stimulation exceeded the termination of the stimulus by about 200ms and is therefore suitable not only for stimulus competition but also for supporting attentional capture (Lev-Ari et al., 2020). Despite this evidence, our understanding of the complete mechanisms and neurophysiological substrates of IOR remains sparse and fragmentary (Dorris et al., 2002).

Here, we tackled this mechanistic question by using a modelling strategy relating RTs to mathematical parameters mimicking the different stages of the neural processes leading to manual or saccadic responses (Carpenter & Williams, 1995; Gold & Shadlen, 2007; Pouget et al.,2009). By exploring the rate of accumulation of evidence toward a decision threshold, our results show that with either manual or saccadic responses, IOR mainly depended on a modulation of the variability of evidence accumulation. The ability of a simplistic stochastic accumulator model to account for IOR effects in a quantitative manner may have important implications. Altered IOR may occur after CNS damage (Bartolomeo et al., 1999; Bourgeois et al., 2012; Sapir et al., 1999; Vivas et al., 2006), in psychiatric conditions (Mushquash et al., 2012), or upon neuromodulation of specific brain areas (Bourgeois et al., 2013b). Examining the underlying parameters of the RT distributions may give novel and interesting insight into these deficits and brain functions rather than simply reporting IOR amplitude or time of occurrence, and can reasonably be expected to be a more direct reflection of the underlying neural processes. In the oculomotor domain, saccade-related neurons in FEF, SC, and Lateral Intraparietal cortex (LIP) are thought to be responsible for sensori-motor transformation (Coles et al., 1997; Everling & Munoz, 2000; Purcell et al., 2012;Schall, 2019), and may therefore be relevant nodes of the IOR network. A strength of our model is its ability to predict IOR behavior. It represents a simple and robust way of conceptualizing IOR in the specific contexts of a target-target paradigm, and of TMS-mediated interference.

Variability between individuals is a common observation in IOR effect in humans and animals and also occurred in the group of human subjects tested here. We found that the variability in individual manual IOR scores was correlated with the Gaussian noise parameter captured by the model. Our findings indicate that individual RT variability in IOR is an important estimate of possible processes taking place during sensory selection. The robustness of modelling approaches depends also on the quantity of data modelled (number of subjects and input trials). This had limited the subtlety of the effects we could study (e.g., collapsing left-sided and right-sided targets conditions together).

Our current results do not support Klein and Redden’s (2018) proposal that “input” forms of IOR occur when the oculomotor system is suppressed (e.g., in manual IOR), and are mainly modulated by the ventral cortical visual stream; “output” forms of IOR may instead be observed when the reflexive oculomotor system is not suppressed (e.g., saccadic IOR), and depend on the activity of the dorsal visual stream. Our present evidence, obtained by modelling the results of a target-target paradigm with either manual or saccadic responses, indicated instead that both manual and saccadic IOR mainly depended on a modulation of the rate of evidence accumulation, whose likely neural correlates implicate fronto-parietal networks important for orienting of attention. In the oculomotor domain, saccade-related neurons in FEF, SC, and Lateral Intraparietal cortex (LIP) are thought to be responsible for the sensori-motor transformation (Coles et al., 1997; Everling &Munoz, 2000; Mushquash et al., 2012; Schall, 2019), and were shown to be relevant nodes of the IOR network (Mirpour et al., 2009, 2019). The involvement of these cortical regions in both saccadic and manual IOR was highlighted in the FORTIOR model (Seidel Malkinson & Bartolomeo,2018), which postulates that both manual and saccadic IOR arise from reverberation of noise enhancing activity within priority maps of the frontoparietal circuit linking FEF and IPS. The noisier saliency output is then read by the manual and saccadic motor systems, leading to IOR. According to the FORTIOR model, differences between the readout capacities of the manual and saccadic effector systems when reading the output of the FEF-IPS circuit may then lead to the dissociations between saccadic and manual IOR. For example, the saccade system seems to be more encapsulated (i.e., less prone to interference and illusions) than the manual response system, and relies on a representation that accumulates visual information and location errors over shorter time windows than the representation used for controlling hand movements (Lisi & Cavanagh,2015, 2017). It is important to note that our model assumes a constant non-decision time, an assumption that might be less accurate for manual responses than for saccadic responses and that different tasks, such as Cue-Target ones, might reveal the contribution of additional parameters to IOR generation (Bompas et al., 2017; Patel et al., 2010; Pouget et al., 2011).

Our present modelling evidence shows that TMS affected manual RTs by operating on the Gaussian noise parameter, similarly to the preTMS IOR effects. However, when trying to explore finer effects by directly modelling the TMS-mediated interference on IOR over the right IPS, we could not identify a parameter that significantly explained this effect. This might result from a lack of power due to the limited number of participants in this experimental group.

The enthusiasm about the neurophysiological approach to track IOR effect is somewhat difficult to reconcile with the small changes in response times found in IOR manipulations (but see Dorris et al., 2002). Yet, to make it truly useful, the approach would ideally benefit from modelling hypotheses that are strong enough to be tested, rejected or validated. Further studies involving neural recordings within parietal and prefrontal brain areas will be required to disentangle the dynamics and network extent of IOR generation.

## Acknowledgements

This work was supported by ANR through ANR-16-CE37-0005 and ANR-10-IAIHU-06.

## Notes

### Competing Interest Statement

The authors have declared no competing interest.

### Summary of Updates

Title, analyses (improved data cleaning and model comparisons) and conclusions. Correction for author information.

